# Anomalous Diffusion as a Descriptive Model of Cell Migration

**DOI:** 10.1101/236356

**Authors:** Igor D. Luzhanskey, John P. MacMunn, Joshua D. Cohen, Lauren E. Barney, Lauren E. Jansen, Alyssa D. Schwartz, Shelly R. Peyton

## Abstract

Appropriately chosen descriptive models of cell migration in biomaterials will allow researchers to characterize and ultimately predict the movement of cells in engineered systems for a variety of applications in tissue engineering. The persistent random walk (PRW) model accurately describes cell migration on two-dimensional (2D) substrates. However, this model inherently cannot describe subdiffusive cell movement, i.e. migration paths in which the root mean square displacement increases more slowly than the square root of the time interval. Subdiffusivity is a common characteristic of cells moving in confined environments, such as three-dimensional (3D) porous scaffolds, hydrogel networks, and *in vivo* tissues. We demonstrate that a generalized anomalous diffusion (AD) model, which uses a simple power law to relate the mean square displacement (MSD) to time, more accurately captures individual cell migration paths across a range of engineered 2D and 3D environments than does the more commonly used PRW model. We used the AD model parameters to distinguish cell movement profiles on substrates with different chemokinetic factors, geometries (2D vs 3D), substrate adhesivities, and compliances. Although the two models performed with equal precision for superdiffusive cells, we suggest a simple AD model, in lieu of PRW, to describe cell trajectories in populations with a significant subdiffusive fraction, such as cells in confined, 3D environments.

## Introduction

Cell migration is integral to a variety of physiological processes including organ development, tissue morphogenesis, wound healing, and immune response. A greater understanding of the motility effects of environmental cues can inform the design of biotechnologies such as movement-directing scaffolds. Research into the relationship between cell migration and cues from the cellular microenvironment increasingly takes advantage of the capability to manipulate properties such as the extracellular matrix (ECM) compliance ^1–7^ and density of cell adhesive ligands ^8–12^.

Descriptive (i.e., empirical) models of migration dynamics facilitate analysis of microenvironment dependence in part by assigning parameters to characterize cells, individually and in aggregate. One of the most commonly used models for describing individual cell migration in 2D is the persistent random walk (PRW) model ^13–15^, whose mathematical formulation was originally developed as modified Brownian motion. Until recently, the migration of adherent cells has been explored almost exclusively on 2D surfaces, but is now investigated in 3D as well, partly due to the advent of bioengineered environments capable of encapsulating cells and more closely capturing *in vivo* conditions ^2,16–19^ Despite its success on 2D surfaces, cell migration is often not well described by the PRW model at any appreciably long time scale in confined 3D environments. Given the increasing use of 3D environments to study cell movement, there is a need for a model that can effectively describe individual cell movement in both 2D and 3D environments. Furthermore, there is a need to be able to predict migration model parameters that vary based on easily quantifiable and controllable extracellular conditions such as growth factor concentration, ECM ligand density and composition, and ECM compliance, all of which are known to have a significant effect on cell migration ^20, 21^.

No simple descriptive model is commonly used to capture a broad range of types of cell motion, from highly constrained to ballistic. We therefore adapted the anomalous diffusion (AD) model for individual and aggregate cell migration. In contrast to normal (free) diffusion, in which the mean squared displacement grows linearly with the time interval *τ*, in anomalous diffusion, the mean squared displacement grows as a power, *τ_α_*, of the time interval, where 0<*α*<2, by definition lending this model the flexibility to describe both sub- and superdiffusive motion. Variants of anomalous diffusion, in which *α* may be constant or *τ*-dependent, accurately describe a variety of physical and biological phenomena ^22–27^; however, there are fewer examples of AD’s use in describing adherent cell migration in the literature ^28–30^, and it has not been used to systematically analyze *individual* cell trajectories to the best of our knowledge.

Given that many cells migrating in 3D are subdiffusive, we undertook to systematically characterize the trajectories of individual cells (and aggregate sample-wide migration) under various extracellular conditions using the AD model. We found that PRW and AD gave similar correlation coefficients for superdiffusive cells, but that the AD model was better at describing subdiffusive cells. The AD parameter *α* more clearly differentiated subdiffusive cells from each other than did the PRW parameter *P* (persistence time). The AD parameters, as well as the PRW parameters were found to predictably vary with geometry, elastic modulus, ECM composition, and ECM ligand density. Therefore, we suggest the AD model is a more robust model of individual cell movement, particularly in constrained, 3D environments.

## Materials and Methods

### Cell Culture

The human breast cancer cell line MDA-MB-231 used for cell migration on 2D coverslips and in 3D gels was a generous gift from Shannon Hughes, MIT. The MDA-MB-231 cells used for motility on 2D hydrogel surfaces were a generous gift from Sallie Schneider at the Pioneer Valley Life Sciences Institute. MDA-MB-231 cells were cultured in Dulbecco’s Modified Eagle Medium supplemented with 10% fetal bovine serum (FBS), 1% penicillin-streptomycin (P/S), 1% L-glutamine, and 1% non-essential amino acids (NEAA, Thermo Fisher Scientific, Waltham, MA). Cells were cultured at 37°C and 5% CO_2_. In the cases were MDA-MB-231 cells were cultured on 2D hydrogel substrates, L-glutamine and NEAA were omitted. Immortalized human mesenchymal stem cells (hTERT) were a generous gift from Linda Griffith, MIT, and were cultured in DMEM supplemented with 10% FBS, 1% P/S, 1% L-glutamine, 1% NEAA and 1% sodium pyruvate. Patient-derived human mesenchymal stem cells were provided by the Texas A&M Institute for Regenerative Medicine and cultured between passage 1 and 5 in alpha Minimal Essential Medium (αMEM), 16.5% FBS, and 3 mM L-glutamine (Thermo). In cases where we created conditioned medium from other cell cultures, cells were seeded at 45,000 cells per cm^2^ in 2.5% αMEM with 3 mM L-glutamine for 72 hours. The medium was removed and filtered through a 0.44μm PES filter (Thermo) before use.

### Preparation of ECM-Coated Coverslips

Coverslips were modified to present peptides and full-length ECM proteins as previously described (^31^ and Fig. 1a). Briefly, glass coverslips were silanized through vapor phase deposition of (3-aminopropyl)triethoxysilane (Sigma-Aldrich, St. Louis, MO, USA), rinsed, functionalized with 10 g L^−1^ *N,N*-disuccinimidyl carbonate (Sigma), rinsed, and functionalized through reactive amines with indicated amounts of RGD or fibronectin or ECM protein cocktails that were inspired by the ECM of tissues as follows: bone: 5 μg cm^−2^ of 99% collagen I and 1% osteopontin; brain: 1 μg cm^−2^ of 50% fibronectin, 25% vitronectin, 20% tenascin C, and 5% laminin; and lung: 2 μg cm^−2^ of 33% laminin, 33% collagen IV, 15% collagen I, 15% fibronectin, and 4% tenascin C (all weight%). Rat-tail collagen I and natural mouse laminin were purchased from Thermo; human tenascin C, human vitronectin, and human osteopontin from R&D Systems (Minneapolis, MN, USA); human collagen IV from Neuromics (Edina, MN, USA); and human plasma fibronectin from EMD Millipore (Billerica, MA, USA). Coverslips were then back-filled with PEG_12_ (Thermo) to block non-specific protein adsorption on any remaining surface area.

**Figure 1:**
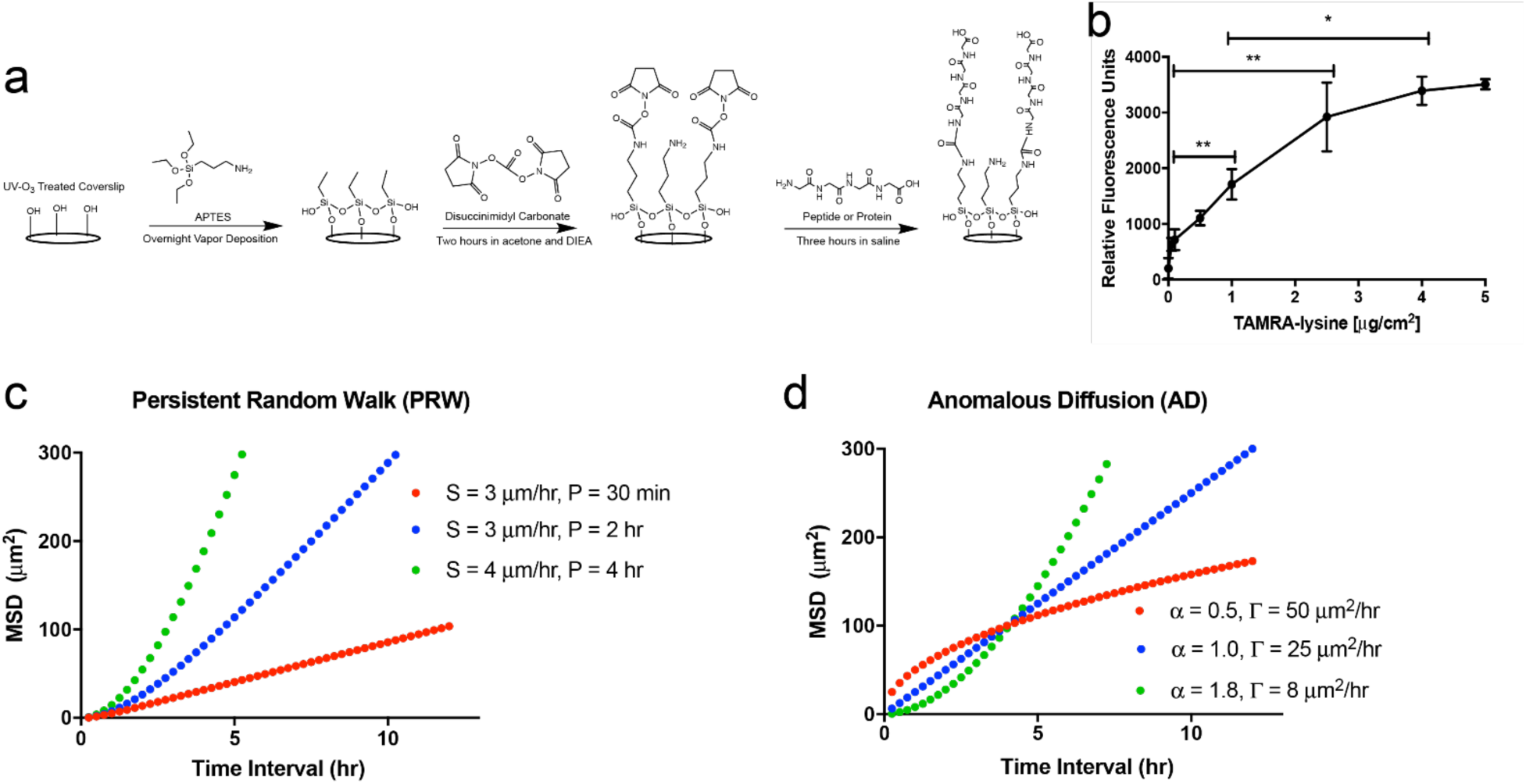
Tunable surfaces and modeling approaches to quantify and describe cell movement. a) Overview of the chemistry used to create ECM-modified coverslips. A three-step process based on silane treatment results in protein- or peptide-modified surfaces (a generic peptide-modified surface is drawn). b) Fluorescence results showing control of peptide surface coupling using the chemistry in (a). Results are from a model peptide (TAMRA-lysine) and read on a fluorometer. c) Theoretical MSD plots for mildly (red), moderately (blue), and highly (green) persistent cell populations (S=speed, and P=persistence time) following the PRW model. d) Theoretical MSD plots for subdiffusive (red), diffusive (blue), and superdiffusive (green) cells following the AD model, each with different, constant (time interval-independent) α and Γ.

A fluorescent assay (Fig. 1b) was used to quantify coupling of a generic, amine-containing peptide (TAMRA-lysine, Anaspec, Fremont, CA) to the coverslip surfaces, as a surrogate for peptides and the proteins used in the reaction depicted in Figure 1. The concentration of TAMRA-lysine was titrated, and fluorescent intensity was measured in a fluorescent plate reader and measured at an excitation/emission of 545/575 nm.

### 2D PEG-PC hydrogels

Glass coverslips (no. 1.5 coverslip glass; Thermo) were plasma treated (Bioforce Nanosciences, Salt Lake City, UT) and subsequently methacrylate-silanized with 2 vol% 3-(trimethoxysilyl) propyl methacrylate (Sigma) in 95% ethanol (adjusted to pH 5.0 with glacial acetic acid) for 2 min, washed 3 times with 100% ethanol, and dried at 40 °C for 30 minutes. PEGDMA (Mn 750, Sigma), from 0.6 to 9.1 wt%, was combined with 17 wt% 2-methacryloyloxyethyl phosphorylcholine (PC) (Sigma) in phosphate buffered saline (PBS). These PEGDMA crosslinker concentrations tune the Young’s moduli of the resulting gels from 0.5 to 4.0 wt% (1 to 64 kPa) ^32^. Solutions were sterilized with a 0.2 μm syringe filter (Thermo) and degassed by nitrogen sparging for 30 seconds. Free-radical polymerization was induced by addition of 0.8 wt% Irgacure (BASF, Florham Park, NJ). Hydrogels of 50 μL per coverslip were covered with an untreated coverslip and polymerized under UV light for 20 minutes. Postpolymerization, hydrogels were allowed to swell for 24 hours in PBS, then treated with 0.3 mg mL^−1^ of sulfo-SANPAH (ProteoChem, Denver, CO; in pH 8.5 HEPES buffer) under UV light for 10 minutes, rinsed twice with PBS, and incubated overnight with 10μg cm^−2^ rat-tail collagen I.

### 3D PEG-Maleimide hydrogels

3D PEG-maleimide (PEG-Mal) hydrogels were prepared at 20 wt% solution with a 20K 4-arm PEG-Mal (Jenkem Technology, Plano, TX) crosslinked at a 1:1 ratio with 1.5K linear PEG-dithiol (Jenkem) in 2mM triethanolamine (pH ~ 7.4). Cells were pelleted and resuspended in PEG-Mal solution before mixing with PEG-dithiol. The cell adhesion peptides (GenScript, Piscataway, NJ) CGP(GPP)_5_GFOGER(GPP)_5_ (1mM), CGPHSRN(G)_6_RGDS (0.8mM), CGGSVVYGLR (0.1mM), and CGGAEIDGIEL (0.1mM) were reacted with PEG-Mal 10 minutes before gelation. Gels were polymerized in 10 μL volumes for 5 minutes before swelling in cell culture medium.

### Cell Migration

On coverslips, cells were seeded at 4000 cells per cm^2^ and given 18 hours to adhere in growth medium. On 2D hydrogels, cells were seeded at 5,700 cells per cm^2^ and given 24 hours to adhere in growth medium. In 3D hydrogels, cells were seeded at 1,000 cells μL^−1^ and allowed to equilibrate and spread within the gels for 24 hours, with a medium change to regular or conditioned medium 4 hours prior to microscopy. On coverslips with ECM-cocktail coatings, seeded MDA-MB-231 cells were treated with a live-cell fluorescent dye (CMFDA, Thermo) and then provided fresh medium or medium supplemented with 40 ng mL^−1^ epidermal growth factor (EGF, R&D Systems) or 0.83 μg mL^−1^ anti-β_1_ integrin (clone P5D2, R&D Systems) 4 hours prior to microscopy. Brightfield and fluorescent images were taken at 15-minute intervals for 12 hours using an EC Plan-Neofluar 10× 0.3 NA air objective (Carl Zeiss AG, Oberkochen, Germany) and cells were tracked using Imaris (Bitplane, St. Paul, MN, USA). Cell migration on 2D hydrogels was done in the absence of fluorescence, and cells were manually tracked. For 3D migration, fluorescent images were taken every 15 minutes for 12 hours with 10 μm z-steps using a Zeiss Cell Observer Spinning Disc at 20X with a NA 0.5 air objective (Zeiss) and cells were tracked using Imaris. *N*≥ 24 individual cell paths were generated for each condition, consisting of coordinates (*x_n_*(*t*), *y_n_*(*t*)) or (*x_n_*(*t*), *y_n_*(*t*), *z_n_*(*t*)) at observation times *t* = 0, Δ*t*, 2Δ*t*, … *t*_max_, where Δ*t* is the time between image acquisitions, *t*_max_ is the last acquisition time, and *n* ranges from 1 to *N*. Cells that contacted other cells, underwent division or apoptosis, or were not fully in frame for the entire 12- or 18-hour observation period were excluded from all calculations.

### Displacement calculations of cell migration paths

The individual mean square displacement (MSD) function 〈*d*^2^〉_*n*_(*τ*) of each cell was calculated for *τ* = 0.25 hr, 0. 50 hr, 0.75 hr … by averaging the square displacements over all available time intervals of length *τ* within the cell’s trajectory (i.e., overlapping intervals ^14^), according to the equation

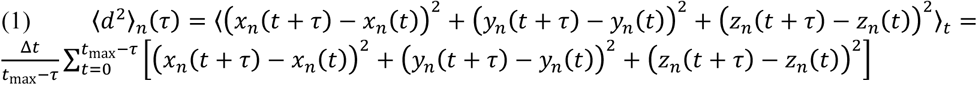

where the angle brackets indicate averaging over all possible starting times *t* and the z-terms were neglected for 2D paths.

The aggregate MSD function 〈*d*^2^〉_agg_(*τ*) of each condition was calculated according to

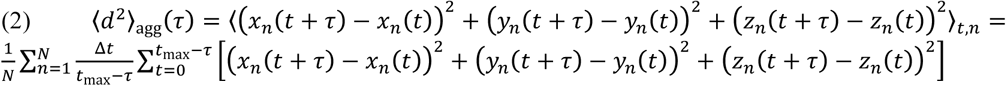

where the angle brackets indicate averaging all available intervals of length *τ* from all available cell trajectories in the condition and the z-terms were neglected for 2D paths.

Fitting of 〈*d*^2^〉_*n*_(*τ*) or 〈*d*^2^〉_agg_(*τ*) to the AD or PRW model functions was performed on points with *τ* values between Δ*t* = 0.25 hr and *τ*_max_ = 4 hr — only up to 1/3 of the maximum, so that each value of 〈*d*^2^〉(*τ*) used in the fitting was obtained by averaging data containing at least 3 statistically independent displacement values.

### The Persistent Random Walk Model

Individual and aggregate MSD data up to a maximum *τ* of 4 hours was fitted to the persistent random walk (PRW) equation used by Dunn ^33^

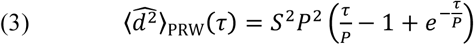

in which *S* is root mean square speed over all available Δ*t*-sized intervals (15 minutes) and *P* is persistence time, a characteristic parameter describing the average length of time over which cells resists major changes in direction of movement. For individual cells, the persistence is *P_n_* and the root mean square speed is calculated as

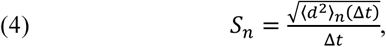

while for cell samples within a given condition, the PRW parameters are *P*_agg_ and

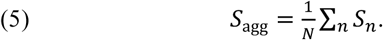

Best-fit values of *P* (*P_n_* for cells and *P*_agg_ for conditions) between 0 and *t*_max_ were obtained using the Matlab (Mathworks) function lsqcurvefit(), which attempts to minimize the residual sum of squares (RSS), defined as

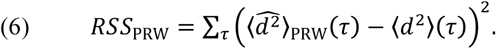

The coefficient of determination (COD), a measure of goodness of fit, was calculated as

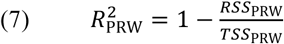

where

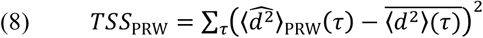

is the total sum of squares (TSS) and the overbar (^_^) indicates the unweighted mean for all *τ* (only up to 4 hours).

Whereas the PRW model has only 1 fitted parameter (P), the AD model has 2. Since additional “degrees of freedom” can be expected to increase the correlation coefficient, some comparisons of the accuracy of the AD and PRW model might be reversed in favor of PRW if both *S* and *P* are allowed to vary. So a 2-parameter PRW fit was performed to obtain additional, arguably more “fair” values of *S, P*, and *R*^2^_prw_. Generally, the use of the 2-parameter fit did not significantly increase *R*^2^_prw_ but sometimes drastically altered the best-fit parameters (see Table S2). Unless otherwise indicated, reported *P* values are from the 1-parameter fit.

### The Anomalous Diffusion Model

The anomalous diffusion (AD) model equation

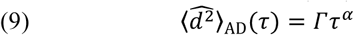

where *α* is the anomalous exponent and Γ is the “anomalous diffusivity” parameter, was linearized by taking the logarithm to obtain

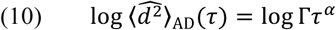

Then log 〈*d*^2^〉(*τ*) *versus* log *τ* data, up to a maximum *τ* of 4 hours, was fitted to the equation using lsqcurvefit() and Microsoft Excel’s slope() and intercept() functions. AD parameters *α*_n_ and Γ_*n*_, for individual-cell MSD data, or *α_agg_* and Γ_agg_, for aggregate MSD data, were simultaneously determined. Best-fit *α* was restricted to between 0 and 2 and best-fit Γ restricted to between 0 and 10000.

For AD model fitting of MSD *vs. τ* data (both individual and aggregate), RSS, TSS, and COD were defined, respectively, as

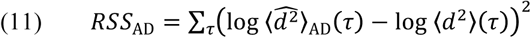

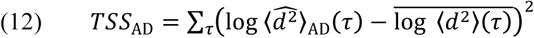

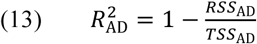

### Data and Model Analysis

A suite of purpose-built Matlab programs was used to process and analyse the migration data at the cell and aggregate level, including model fittings, and report, classify, and compare the results, and produce analytical plots such as Γ-binned-by-α plots and histograms. The MSD calculations and PRW model fittings were confirmed using the Visual Basic program DiPer ^34^ The mean, median, quartiles, 10^th^, and 90^th^ percentiles of individual-cell best-fit parameters were determined for each condition, and GraphPad Prism and Matlab were used to make the plots. Cell-specific parameters such as *α_n_* and *P_n_* were not weighted differently depending on the strength of the fit, i.e. *R*^2^ value, when calculating condition-wide averages. Individual cells were considered subdiffusive for descriptive purposes if 0 ≤ *α_n_* ≤ 1 and superdiffusive if 1 < *α_n_* ≤ 2.

### Ethics Approval

Ethics approval is not required for this study. Conditioned medium from patient cells described in Figure 5 were de-identified and exempt from IRB approval.

## Results

### The AD model outperforms PRW in describing individual subdiffusive cell motion

We first quantified cell motility on supra-physiologically stiff surfaces: 2D coverslips coupled with full-length, integrin-binding (ECM) proteins. We created three different surfaces, inspired by proteins found in different tissues of the human body: bone, brain, and lung (Fig 2). Independently, we perturbed MDA-MB-231 chemokinesis and adhesivity, chemically, by adding either EGF (green) or a function-affecting antibody to β_1_ integrin (red) (Fig. 2a-c). On these rigid surfaces, regardless of the ECM protein cocktail or chemical perturbation, cells were largely (28-84%) superdiffusive (1<*α_n_*<2, Fig. 2d-f, Table S.1). Most cells were well described by the AD model, with individual 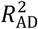≥0.8 for >90% of cells in this experiment and this study (Table S1 and Fig. S1).

**Figure 2:**
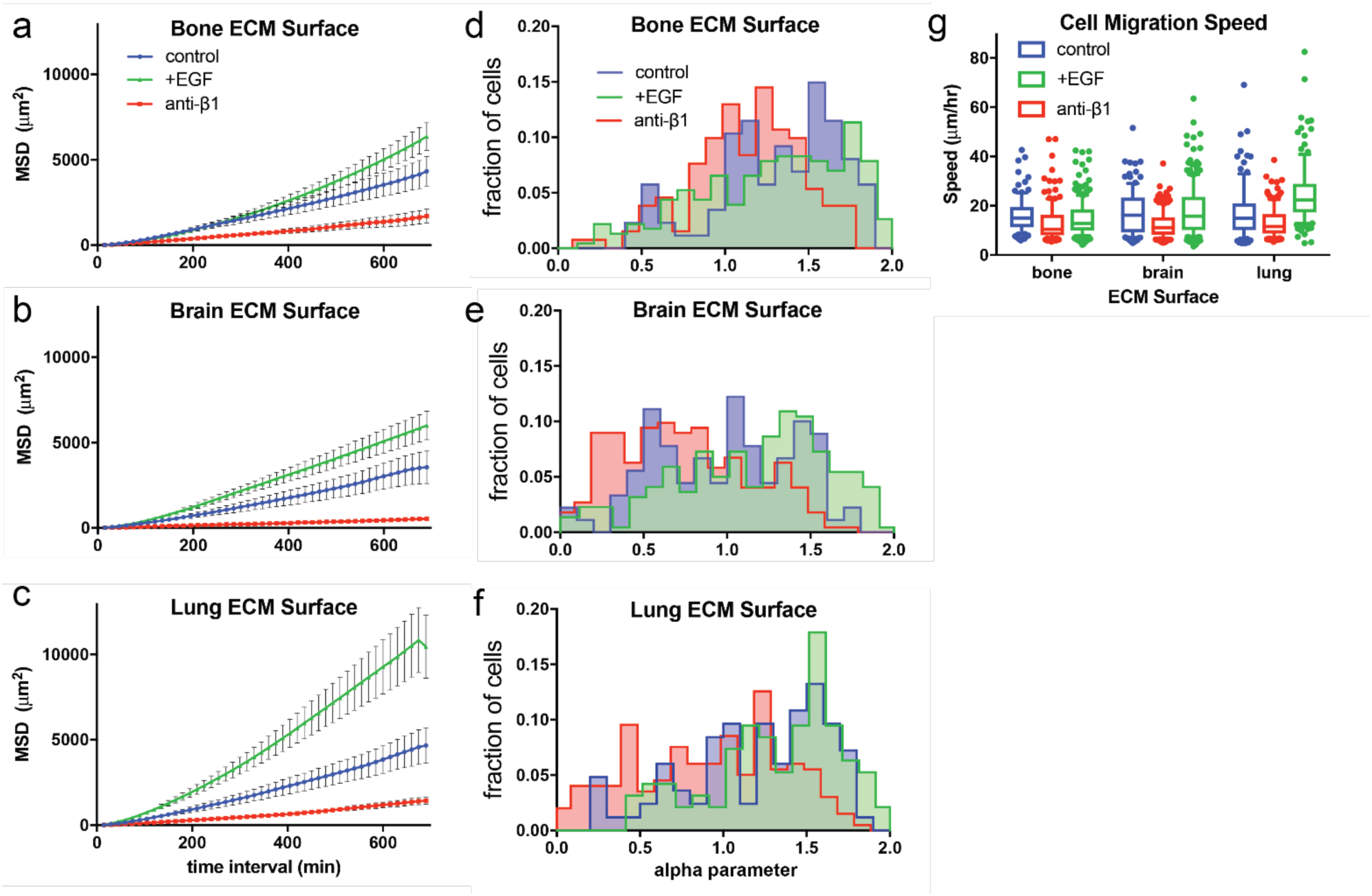
The alpha parameter of the anomalous diffusion model is highly sensitive to chemokinetic perturbation. The mean squared displacement is shown for all cells and time intervals (a-c), and best-fit estimates for α_n_ as histograms (d-f), for the different ECM surfaces created: bone (a, d), brain (b, e), and lung (c, f). (g) Individual average cell speed box-and-whisker plots for all 9 substrate and treatment conditions. Control experiments (performed in standard growth medium) are shown in blue, supplemented with EGF (green), and treated with a function-affecting antibody to β_1_ integrin (red).

Across all conditions, both 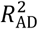 and 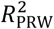 approached 1 as *α_n_* approached its maximum of 2 (Fig. S1). Given the flexibility of fitting for PRW, and that both models fit well, this is an argument for using PRW for cells on rigid 2D surfaces. While individual 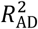 and 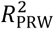 remained greater than 0.95 for 97% of superdiffusive cells (*α_n_*>1), 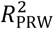 decreased significantly as *α_n_* decreased below 1 (subdiffusive cells, Fig. S1). 82% of subdiffusive cells had 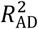>0.8, while 45% of subdiffusive cells had 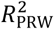>0.8, indicating that the AD model fit cell MSD data far better than the PRW model. Therefore, when considering all cells from each population, the AD model outperformed the PRW model across all ECMs (Table S.1). Finally, fitting with the AD model on an individual cell basis revealed population heterogeneity (in terms of the distribution of *α* and Γ within a population compared to the distribution of *P* and *S*) that persisted over the tracked time interval, which would have been missed by analysis with PRW alone (Table S.1-S.3, Fig. S1).

### The anomalous exponent, α, is most sensitive to integrin-binding to 2D rigid substrates

We observed sub- and super-diffusive behavior in individual cells during 2D migration on bone-, brain-, and lung-inspired-coated glass coverslips (Fig. 2). This is in line with previous studies of mammalian cell migration on other hard tractable ECM-coated surfaces ^35^, which have previously been described with the PRW model. Cells exhibited much longer displacements on the lung ECM compared to brain and bone surfaces (Fig. 2a-c). Blocking β_1_ integrin suppressed migration, while EGF treatment enhanced it on all surfaces, and lung-like surfaces tended to have greater migration-enhancing abilities than brain- and bone-like surfaces. When comparing the effects of these soluble factors versus the cell-adhesive proteins on the surfaces, EGF and β_1_-integrin targeting had a greater effect on the MSD, and on the AD model parameters *α* and Γ, than surface ECM.

The *α_n_* distribution within each condition was typically unimodal and sensitive to the ECM adhesivity and soluble factors, highlighting the capability of the power-function model to describe a heterogeneous population of cells (Fig. 2d-f, S2). Regardless of the ECM protein cocktail or chemical perturbation, cells’ individual anomalous exponents spanned the entire possible range 0 – 2 but tended to have a majority of superdiffusive cells, with superdiffusive fraction ranging from 28% on brain ECM-like surface with anti-ß_1_ integrin to 84% on bone ECM-like surface with no chemical perturbation (average 63%; Table S.1). Furthermore, all 9 conditions had an aggregate anomalous exponent greater than 1, indicating aggregate superdiffusive movement, with *α*_agg_ ranging from 1.02 to 1.58 (average 1.40, Table S3).

The distribution of both *α_n_* and Γ_n_ shifted in response to soluble factor treatment, while *α_n_* was more sensitive to substrate variation than Γ_n_ (compare Fig. 2d-f with S2). Average cell speed was more sensitive to chemical perturbation than substrate adhesivity (Fig. 2g). Treatment with anti-β_1_ integrin decreased median *α*, cell speed, and MSD across all three substrates, while EGF treatment increased median *α* and average displacement. Overall, for sample populations where both PRW and AD fit well, α increases with cell speed and persistence time, while there were no observed trends with Γ. Across all cell data, as expected, persistence time and α strongly positively correlated (Fig. S3), as did Γ and speed.

On average, a greater fraction of cells were superdiffusive in EGF-treated conditions (73%) than in untreated (69%) or anti-ß_1_-treated conditions (48%). To fairly compare the effect on the anomalous diffusion coefficient *Γ*_n_ of different stimulants, we segregated individual cells by α_n_ so that individual values of Γ_n_ would have approximately the same units (e.g. from μm^2^/hr^1.6^ to μm^2^/hr^2.0^). When segregated in this way, Γ_n_, much like α_n_, was higher for EGF-treated cells than for untreated cells and anti-β_1_-treated cells; and higher for cells on lung-like than on brain- or bone-like ECM cocktail (Fig. S.3).

### Γ, not α, is sensitive to substrate adhesivity

Given the known dependence of cell migration speed on the density and type of adhesive surface ligands ^35, 36^, we examined the quality of the AD model, in comparison to the PRW model, on surfaces of varying densities of adhesive ligand. We created coverslip surfaces coupled with either the integrin-binding peptide RGD (Fig. 3a, c, e, and g) or full length ECM protein fibronectin (Fig. 3b, d, f, and h) using the same silane coverslip chemistry depicted in Figure 1 and applied in Fig. 2. To determine if the results from Figure 2 were limited to the epithelial breast cancer cell line profiled, we expanded the study to a bone marrow-derived mesenchymal stem cell line (MSC). Virtually all cells were superdiffusive (Table S1 and Fig. 3e-f), with a much narrower *α_n_* range than observed for 231s on the 2D tissue-specific ECM-mimicking surfaces.

**Figure 3.**
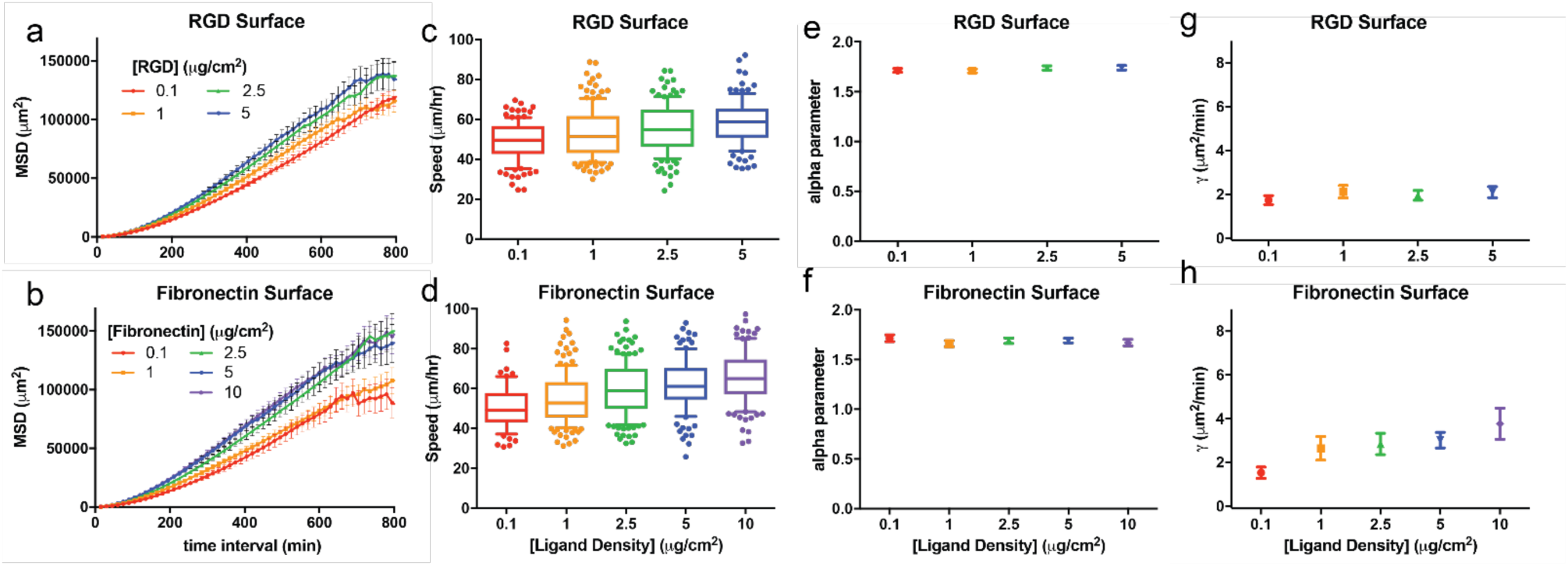
Gamma follows MSD and cell speeds during haptokinesis. Average MSD versus time intervals for hTERT MSC cell migration on surfaces functionalized with different concentrations of RGD (a) or fibronectin (e). Individual cell speed box-and-whisker plots are shown for RGD (c) and fibronectin (d) surfaces. (e-h) The anomalous diffusion parameters α_n_ (e-f) and Γ_n_ (g-h) are given for RGD (e,g) and fibronectin (f,h) surfaces.

For both adhesive ligands used, aggregate MSD increased with ligand concentration (Fig. 3a-b). Median cell speed (Fig. 3c-d) and median Γ_n_ (Fig. 3g-h and S4) increased with increasing fibronectin concentration, while median α_n_ (Fig. 3e-f and S4 and Table S1) and median persistence time *P_n_* (Table S2) showed no observable dependence on surface ligand concentration. Median α_n_ was 1.73 to 1.76 for cells on RGD, and 1.71 to 1.73 for cells on fibronectin. These results suggest that greater ligand density increased random cell motility, and thus increased speed and Γ, with small changes to MSD. However, there was no effect on cell directionality, which explains why we observed no differences in α nor persistence time. Regardless, due to the largely superdiffusive cell populations for all conditions tested, average individual R^2^ for both the PRW and AD models for all samples in this experiment was at least 0.98 (Table S1).

### Γ and speed have a biphasic dependence on substrate stiffness

Given the known dependence of cell migration speed on substrate stiffness ^1,37^, we tested the AD model against cell migration data obtained from MDA-MB-231 breast cancer cells on substrates of varying stiffness. We used our previously published PEG-PC hydrogel system, which has independent control over modulus and the density of adhesive ligands on the substrate surface ^32^. We varied the substrate modulus from 1 to 64 kPa, and coupled type 1 collagen to the surface to make it cell adhesive (Fig. 4). The MSD of the breast cancer cells had a biphasic dependence on substrate stiffness, with maximum MSD occurring on 18 kPa gels (Fig. 4a), and similar to previous reports on biomaterial surfaces, we observed a biphasic dependence of cell migration speed as a function of gel stiffness (Fig. 4b)^1^. Somewhat in parallel with this biphasic response, *α_n_* and *S_n_* slightly positively correlated among cells on low- and high-modulus surfaces and slightly negatively correlated among cells on medium-modulus surfaces (Fig. S6).

**Figure 4:**
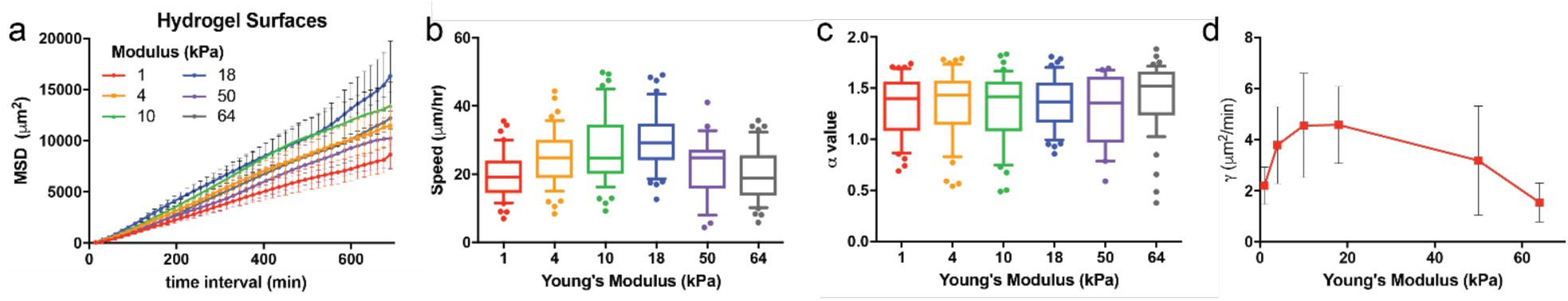
Cell mean squared displacement, speed, and the diffusion coefficient are sensitive to substrate modulus. Average MSD versus time intervals for breast cancer cell migration on hydrogel surfaces coupled with a constant density of collagen (10μg cm^−2^) and varying substrate modulus (1-64 kPa). Individual cell speed box-and-whisker plots are shown each stiffness tested. (c-d) The anomalous diffusion parameters *α_n_* (c) and Γ_*n*_ (d) are given for each stiffness tested.

Most cells migrating on these 2D soft gels were superdiffusive, with *α* values averaging between 1.3-1.4, and no observable relationship with gel modulus (Fig. 4c and S5b). Instead, there was a visible correlation between cell speed and Γ, and were both maximized on the 18 kPa surface (Fig. S2, Tables S2-3, and Fig. 4d). We conclude, as in Fig. 3, that the Γ parameter (acting as a transport diffusivity) is associated with cell speed, and the α parameter, (the trajectory anomality^38^ is not. Furthermore, cell migration was superdiffusive regardless of gel stiffness (Fig. 4d, S6a), or ligand density (Fig 3c,g), for the range of conditions we tested. For all samples, the average individual *R*^2^_AD_ was at least 0.95 and the average individual *R*^2^_PRW_ was at least 0.77 (Table S1).

### Cell migration in confined, 3D environments is largely subdiffusive

Cell motility models, such as PRW, were initially developed from data obtained from cells migrating on flat, 2D surfaces. However, *in vivo*, cell movement is largely in 3D, with cells surrounded by ECM and other cells. To test the effectiveness of the AD model in describing this type of confined cell movement, we used another PEG-based gel ^39–41^ and measured cell motility in 3D as we exposed them to different pro-migratory chemical stimulations (conditioned medium from patient cell cultures). The gels were crosslinked with MMP-sensitive peptides, but the mesh size was orders of magnitude smaller than a cell (~20-25nm). Therefore, cell movement in this environment is confined, and cells are expected to exhibit subdiffusive motion.

We observed that all chemical stimulations increased the displacement of cells in 3D compared to normal growth medium (Fig. 5a), although speed across the population was not dramatically affected. Mean speeds of cell populations slightly increased as a function of stimulation (Table S.2), and individual cells were found to be faster in conditions with medium from either Patient 1 or Patient 3 compared to all cells in normal growth medium (Fig. 5b). 3D migration is largely subdiffusive (*α* < 1) in control medium and is approximately diffusive (*α* ≈ 1) in MSC-conditioned medium (Fig. 5c). *α* increased significantly for cells in Patient 1 and Patient 2 MSC-conditioned medium compared to control medium.

**Figure 5:**
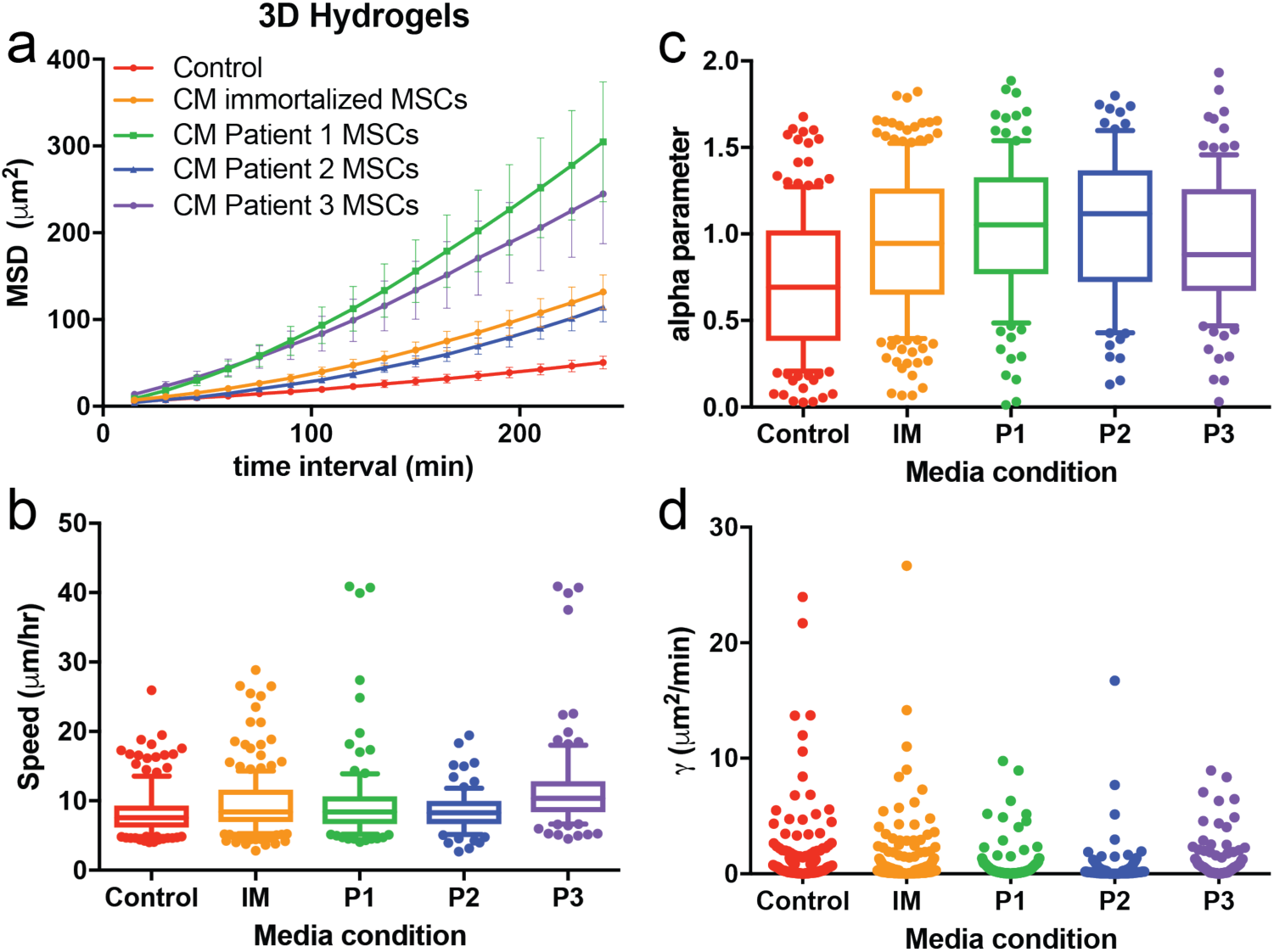
Alpha describes subdiffusion of cells in soft, 3D hydrogels. a) MSD as a function of time interval for breast cancer cells migrating in soft, 3D hydrogels supplemented with either normal growth medium (control), or conditioned medium (CM) from different sources of MSCs (Immortalized MSCs: IM, Patient 1: P1, Patient 2: P2, Patient 3: P3). b) Individual cell speed box-and-whisker plots for the different medium conditions. (c-d) The anomalous diffusion parameters α_n_ (c) and Γ_n_ (d) are given for each medium condition tested.

The Γ parameter had low values overall compared to the cells in 2D environments (Figs. 2-5) and was less affected by these medium conditions, revealing a stronger relationship with dimensionality compared to medium (Fig. 5d). As shown in Table S.1, the AD model fits exceeded the PRW fits for all conditions. AD and PRW fit equally well (around 0.95, Table S.1) for superdiffusive cells, but the goodness of fit for PRW rapidly dropped off as α decreased below 0.8 (data not shown). We conclude that the PRW model is not applicable for cells in 3D environments because of the very high proportion of subdiffusive cells. The AD model, as a power-function model, is the simplest adjustable monotonically increasing scale-invariant function, and is sufficient to capture the heterogeneity of these cell trajectories.

## Discussion

By analyzing different human cell lines in unique engineered biomaterial environments, we demonstrate that, overall, the AD model describes cell movement better than the commonly used PRW model. The exception to this rule is best shown in the experiments from Figures 3 and 4, which contained the highest proportion of superdiffusive cells. Superdiffusive motion is described equally well by both models, with PRW arguably giving more readily interpretable information (namely persistence time and the relation between persistence time and adhesivity or elasticity). However, the AD model was equally good at relating motility behaviors to its own model parameters. The largest advantage of using the AD model was in 3D environments, wherein the largest population of cells were subdiffusive.

Comparisons between the 2D (Fig. 2) and 3D (Fig. 5) experiments highlighted the divergence in fitting between these two models, because these were the conditions that contained the highest proportion of subdiffusive cells. PRW failed to describe the motion of cells in confined, 3D environments. In these cases, the AD model fit exceeded that of PRW. In 3D, where subdiffusive movement is expected, the PRW equation often yields a poor fit, with P≈0 such that either the trajectory is diffusive with speed *S* (the MSD function is linear) or the fitted parameters do not carry meaning. Cells in 3D confined environments have MSD functions unlimited to diffusive or superdiffusive paths, and thus need to be described by a model with flexible parameters, such as AD.

The effect of the environment on a cell’s ability to move away from previous locations at a slower rate than if it “freely diffused” is reflected in the value of *α*. Although not explored here, high persistence, and a strong correlation between cell speed and persistence, are observed in both 2D free diffusion and for 1D-confined paths ^42^ For diffusive and superdiffusive cells, P varies positively with α, which was generally increased by EGF and by MSC-conditioned medium, and decreased by anti-β1 integrin antibody. Targeting integrin β_1_ increased the population of subdiffusive cells, supporting the known role for integrins as critical for polarization and persistent motion ^43^. The variation in the integrin-binding domains presented in different ECM cocktails also affected the MSD of cells, but treatment with EGF had the most significant impact on increasing the proportion of superdiffusive cells (Fig. 2). This is not surprising given the chemo-kinetic potency of EGF ^44^.

The range of *α* for cells in the 3D gels spanned from very subdiffusive (practically immobile) to very superdiffusive (ballistic motion, equivalent to moving in a straight line). *α* increased significantly for cells in Patient 1 and Patient 2 MSC-conditioned medium compared to control medium. Also, both aggregate and α_n_-grouped average individual Γ increased for Patient 1 and Patient 3 MSC-conditioned medium compared to control medium (Table S3, Fig. S7). The AD parameters α_n_, Γ_*n*_, α_agg_, Γ_agg_ similarly increased in EGF-treated medium and decreased in anti-ß1 conditions (Table S1, S3; Fig. 2d-f, S3). Thus Γ and *α* are each independently responsive to chemical perturbation. On the other hand, *α* does not strongly depend on substrate adhesivity (Fig. 3e-f) or elasticity (Fig. 4c) while Γ does (Fig. 3h, 4d).

In this study, the AD MSD equation describes cell migration paths with excellent fit, and contains parameters that clearly depend on the biomaterial or growth factor condition. While our study highlights the robustness of AD to describe cell movement, it is limited to using AD as a descriptive model and not a predictive one. In order to make a predictive model, future studies are needed to understand how α varies with experimental conditions, how Γ is tied to *α*, and how Γ varies with experimental conditions independently of *α*. The dependence of the AD parameters on substrate properties, growth factors, chemokines, and cytokines should be studied further to determine precise predictive correlations. Especially important are experiments with permeable 3D substrates to test the effects of varying elasticity and ligand density on subdiffusive movement. This would confirm that Γ is a strong proxy for cell motility in the absence of obstruction and not simply a reflection/artifact of high *α*.

A disadvantage of the AD model is that the parameters do not have readily apparent physical interpretations. We saw that the average value of Γ, which typically followed closely with cell speed, increases with increasing ECM ligand density and is biphasic with respect to elasticity. If high ligand density and a mid-range/optimal elastic modulus help the cell attain maximum traction, perhaps Γ is a reflection of its magnitude.

It should be noted that we observed a very strong correlation between α and P for all conditions (Fig. S3). Furthermore, Γ_*n*_ strongly correlated with *S_n_* for cells with similar α_n_ (data not shown). Together, these results suggest that the pairs (α_n_, Γ_*n*_) and (*P_n_*, *S_n_*) contain similar information. Nevertheless, the AD model more conveniently represents data with a large fraction of subdiffusive cells because of the mathematical flexibility of *α*. The AD model thus has a broader “dynamic range” with respect to cell motility patterns and arguably reveals rather than masks heterogeneity within populations of subdiffusive cells.

Overall, our study highlighted the increased flexibility, and therefore better fit of the AD model compared to the more commonly used PRW model, particularly in instances for subdiffusive motion, such as in 3D environments. Overall we found that *α* was particularly sensitive to chemical stimulation (soluble factors in the medium or integrin-inhibition), and Γ was more sensitive to substrate adhesivity and elasticity, tracking with cell speed for both. We recommend that the AD model can serve as a basis for simple computational models of cell speed in all environments, with specific applications in tumor growth, wound healing, or the colonization of an artificial tissue engineering scaffold seeded with cells.

**Supplementary Material**

See the supplementary material for complete data for cell motility parameters and model fitting data.

## Acknowledgements

This work was funded by an NIH New Innovator award (DP2CA186573), a CAREER award from the NSF (DMR-1454806), and an S&T award from the University of Massachusetts awarded to SRP. LEB was partially supported by T32 GM008515 from the National Institutes of Health and ADS was supported by a National Science Foundation Graduate Research Fellowship (1451512). SRP is a Pew Biomedical Scholar supported by the Pew Charitable Trusts and a Barry and Afsaneh Siadat Career Development Faculty fellow.

